# A non-canonical role for Jagged1 in endothelial mechanotransduction

**DOI:** 10.1101/2024.11.14.623558

**Authors:** Freddy Suarez Rodriguez, Noora Virtanen, Elmeri Kiviluoto, Rob C. H. Driessen, Feihu Zhao, Carlijn V. C. Bouten, Oscar M. J. A. Stassen, Cecilia M. Sahlgren

## Abstract

The Notch signaling pathway plays a crucial role in regulating endothelial biology. Notch signaling is sensitive to hemodynamic forces and governs mechanically-driven cardiovascular development, physiology, and remodeling. However, the mechanisms by which mechanical forces integrate with the Notch pathway remain largely unknown. Here, we uncover a non-canonical role for the Notch ligand Jagged1 in regulating the activity of mechanosensitive kinases in endothelial cells. We show that stress induces expression and relocalization of Jagged1 to cell junctions downstream of flow. Jagged1 expression under stress demonstrates magnitude dependence and peaks at 0.8-1Pa without impacting Jagged1s Notch-activation potential. On the contrary Jagged1 regulates the activity of mechanosensitive kinases. Deletion of Jagged1 reduces the activity of VEGFR2 and ERK in vitro and diminished ERK activity in zebrafish embryos without affecting canonical Notch signaling. Furthermore, the direct physical stimulation of Jagged1 using antibody-conjugated beads triggers the activation of VEGFR2 and ERK, mediated by Jagged1-induces Src activation. Taken together, we demonstrate a novel non-canonical role for Jagged1 as a regulator of the activity of pathways involved in endothelial mechanotransduction.

## INTRODUCTION

Mechanical forces generated by blood flow direct cardiovascular morphogenesis, homeostasis, and disease progression (Baeyens et al., 2016; Roux et al., 2020). Endothelial cells lining the lumen of cardiovascular tissues are exposed to fluid shear stress (FSS), which is an essential biophysical signal for vasculogenesis, angiogenesis, arterial specification, vessel maturation, and barrier function (Baeyens et al., 2016; Li et al., 2019; Souilhol et al., 2020). FSS is also a well-known predictor of atherosclerotic plaque formation, with zones of lower magnitude of FSS and higher oscillatory shear index (OSI) being atheroprone, and those with laminar-like (low OSI) and higher magnitudes of FSS considered atheroprotective (Baeyens et al., 2016; Souilhol et al., 2020; Tamargo et al., 2023). Physiological FSS varies in different vessels, typically ranging from magnitudes as low as 0.4 Pa to as high as 7 Pa (Baeyens et al., 2016; Katoh, 2023; van Haaften et al., 2017). Endothelial cells express mechanosensors such as the primary cilium, the glycocalyx, Piezo ion channels, the VEGFR2 - VE-cadherin - PECAM junctional mechanosensory complex, and PlexinD1, which activate downstream kinases including ERK, AKT and Src, regulating endothelial cell responses to FSS (Aitken et al., 2023; Rahaman et al., 2023; Tanaka et al., 2021). Src family kinase occupy a special role in FSS signaling by being activated both upstream and downstream of VEGFR2, aiding in initiating and mantaining signaling activation (Aitken et al., 2023; Jin et al., 2003; Miller and Sewell-Loftin, 2022).

The Notch signaling pathway is an essential regulator of cardiovascular development and homeostasis (Cavallero et al., 2021; Fernández-Chacón et al., 2021; MacGrogan et al., 2018). Notch plays an important role in angiogenesis, arterial specification, endothelial barrier function, cardiac morphogenesis, and arterial remodeling (Fernández-Chacón et al., 2021; Grego-Bessa et al., 2007; Hasan and Fischer, 2022; Polacheck et al., 2017; Priya et al., 2020; Tian et al., 2017). In canonical Notch signaling, Notch ligands on the membrane of signal-sending cells bind and activate Notch receptors on signal-receiving cells. Activation of the receptor requires endocytosis of the ligand after receptor binding (Seib and Klein, 2021). The activated receptor is then cleaved, and its intracellular domain is translocated to the nucleus, where it promotes the transcription of Notch target genes (Falo-Sanjuan and Bray, 2020). Notch activity is mechanosensitive, and Notch signaling is essential for the cellular response to hemodynamic stress (Cavallero et al., 2021; Stassen et al., 2020; Suarez Rodriguez et al., 2023; Tsata and Beis, 2020). Laminar-like and high magnitude FSS, above 2 Pa, has been shown to enhance Notch1 activity (Fang et al., 2017; Mack et al., 2017), while OSI of high (1.5 Pa) and low (0.4 Pa) magnitudes has been found to stimulate Notch3 and Notch4 activity, respectively (Karthika et al., 2023; Souilhol et al., 2022).

FSS-induced Notch1 activity in arteries promotes vascular barrier integrity (Polacheck et al., 2017) and serves an atheroprotective role in regions exposed to high laminar blood flow (Mack et al., 2017). In contrast to Notch1, recent data suggest that the Notch ligand Jagged1 (Jag1) is enriched in pro-atherogenic zones, of FSS with low magnitude and high OSI (Souilhol et al., 2022; Sreelakshmi et al., 2024), and promotes atherosclerosis (Souilhol et al., 2022). However, the exact role of Jag1 in the endothelium remains unclear. Here, we set out to investigated the flow-responsive nature of Jag1, and its role in endothelial mechanotransduction. We show that magnitude-specific changes in FSS tune Jag1 expression and localization and identify a non-canonical role of Jag1 in regulating the activity of FSS-responsive kinases.

## RESULTS AND DISCUSSION

### Shear stress induces changes in Jagged1 expression and localization inconsistent with canonical Notch activation

We have previously demonstrated that FSS induces clustering of Jag1 into submembranous vesicles, enhances Jag1 endocytosis and Jag1-induced Notch activation in VSMCs and reporter cells adjacent to the endothelium (Driessen et al., 2018; van Engeland et al., 2019). Previous data have shown that Notch1, the main receptor expressed in endothelial cells (Briot et al., 2015), polarizes in the direction of flow (Giese et al., 2025; Mack et al., 2017). Here we analysed the relocalization of Jag1 in response to flow. Surprisingly, in addition to the contralateral localization needed for Jag1-Notch transactivation, FSS also induced a signficant Jag1 polarization in the direction of flow in both human aortic endothelial cells (HAoECs) (Fig. 1A-C) and human umbilical vein endothelial cells (HUVEC) (fig. S1A-C) exposed to ∼1Pa for 24 hours.

**Fig. 1.**
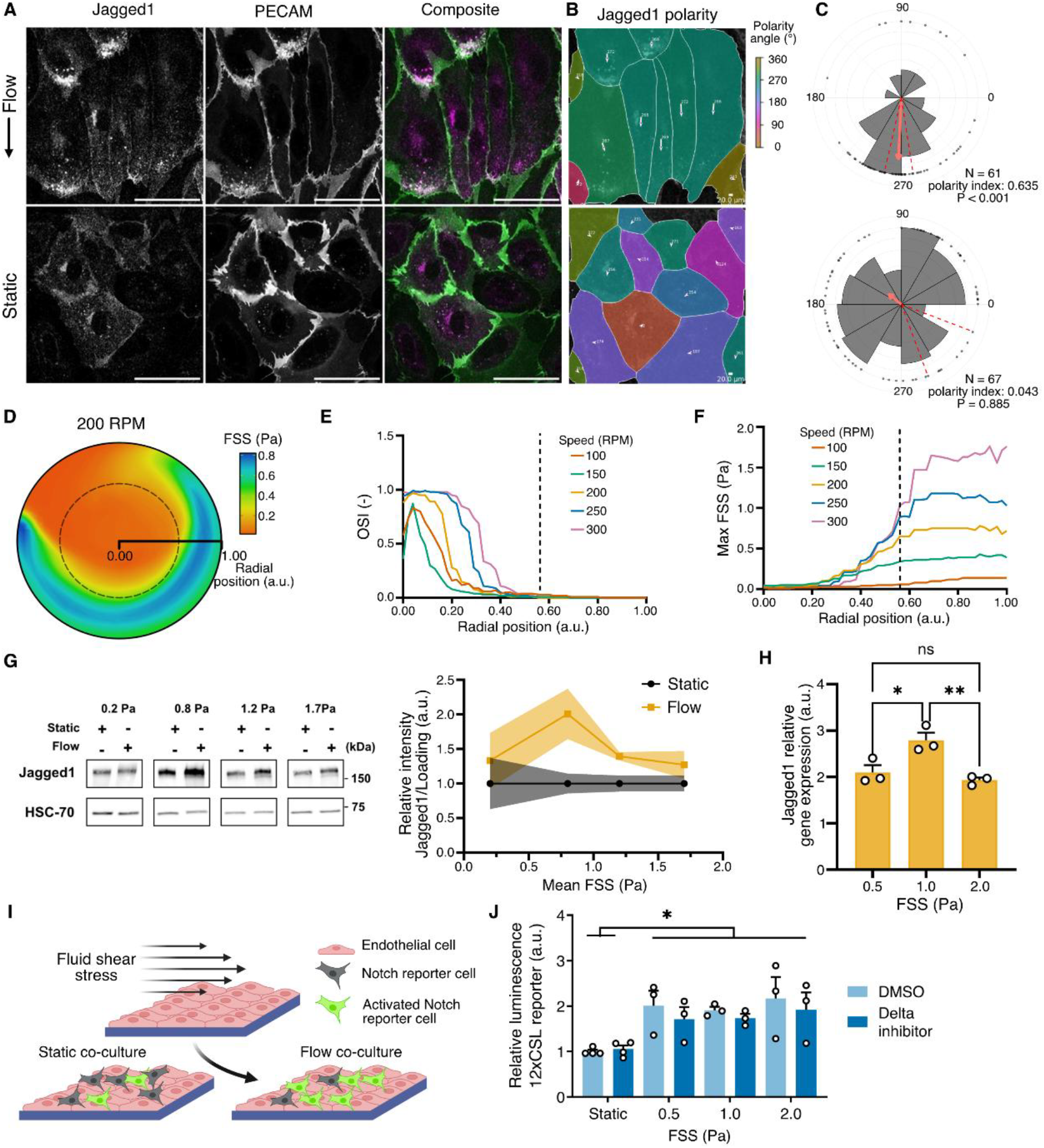
Jagged1 expression and localization in response to FSS is decoupled from its transactivation potential. (**A**) Confocal microscopy images of HAoECs exposed to ∼1 Pa laminar and continuous FSS for 24 hours in Ibidi® microfluidic chips. Jag1 (magenta) demonstrated polarized localization in the direction of flow. PECAM/CD31 (green) was used to denote cell junctions. (**B**) Analysis of Jag1 polarization using PolarityJaM Python API. Mask color represents polarity angle. Vector size and orientation represent magnitude and angle of the polarization. (**C**) Merged graph and analysis of independent experiments in PolarityJaM web app. Vector represents the circular mean direction, doted lines represent the 95% confidence interval. Rayleigh test of uniformity was used to analyze statistical significance. (**D**) Computational fluid dynamic analysis of the FSS pattern in an orbital shaker system using 6-well plates. The radial position indicates the radial coordinates of the well, with 0.00 indicating the center of the well and 1.00 the wall of the well. The dotted line separates the wells’ outer area with high magnitude laminar flow from the inner area with highly oscillatory flow. Cells were collected from the outer area of the well, from radial position 0.56 to 1.00. (**E**) Oscillatory shear index (OSI) was calculated from the simulation and used to determine the zones with the lowest oscillatory flow. (**F**) Maximum FSS levels were used to determine the magnitude of FSS at each speed (RPM). (**G**) Western blot (WB) analysis of Jag1 protein levels in HUVECs exposed to different magnitudes of FSS using our orbital shaker system. The values are presented relative to the corresponding static control. (**H**) Q-PCR analysis of Jag1 gene expression levels in HUVECs exposed to laminar and continuous FSS of different magnitudes for 24 hours in Ibidi® microfluidic chips. (**I**) Schematic of experiment, Notch reporter cells cultured on HUVECs exposed to shear. (**J**) Notch activity in 12xCSL-Luciferase expressing HEK293-FLN1 cells co-cultured with HUVECs exposed to different magnitudes of FSS. Delta signaling was inhibited by the fucose analog 6-alkenyl-fucose. All experiments were performed three times, for H this included three technical replicates within each experiment. The levels are presented as the mean of each replicate relative to their corresponding control + standard error of the mean (SEM). For WBs HSC-70 was used as a loading control. P-values were obtained with GraphPad Prism as described in the methodology. Significance is indicated as: ns p > 0.05, * p < 0.05, ** p < 0.01, *** p < 0.001. Arbitrary units are indicated as (a.u.).

Notably, while Jag1 expression and Notch1 activity increase in cells exposed to FSS (Chen et al., 2024; Fang et al., 2017; Mack et al., 2017; Souilhol et al., 2022; van Engeland et al., 2019), their response to changes in FSS magnitude differs. Notch activation has been found to increase with the magnitude of the FSS beyond 2 Pa (Fang et al., 2017; Mack et al., 2017). In contrast, Jag1 expression was reported to be the highest at low magnitudes of FSS (0.4 Pa) (Souilhol et al., 2022).Given the importance of FSS magnitude, we next evaluated the effect shear stress magnitude on the expression of Jag1 and other components of the Notch pathway. To this end, we used an orbital shaker to complement the microfluidic chips and obtain sufficient material for further biochemical analysis. We utilized computational fluidic dynamics to characterize FSS magnitude and OSI in a 6-well cell culture plate (Fig. 1D) to be able to isolate the cells exposed to laminar-like flow with low OSI (Fig. 1E) and a specific magnitude of FSS (Fig. 1F). Using this system we confirmed that FSS induces the highest Jag1 expression at low magnitudes of FSS, with protein levels decreasing at 1.2 Pa and beyond (Fig. 1G) in line with what has been reported (Souilhol et al., 2022). Additionally, we reveal that protein levels initially peak at approximately 0.8 Pa before decreasing. To assess if FSS affected Jag1 gene expression, we analysed mRNA levels of the main ligands and receptors expressed in endothelial cells, Jag1, Dll4, Notch1 and Notch4 through qPCR of cells flowed using Ibidi® microfluidic chips. Jag1 expression peaked at 1 Pa (Fig 1H), further corroborating our results. On the contrary, the expression of Notch receptors did not change significantly at the magnitudes evaluated (fig. S2).

The changes in Jag1 expression below 2 Pa (Fig. 1G-H) (Souilhol et al., 2022) are in discrepancy with the continuous increase in Notch activity reported by the literature for FSS above 2 Pa(Fang et al., 2017; Mack et al., 2017). We next assessed how different FSS magnitudes influenced Jag1 signaling potential by co-coculturing endothelial cells exposed to FSS together with Notch reporter cells (Fig. 1I) (Driessen et al., 2018). As endothelial cells also express Dll4, we inhibited Delta signaling using fucose analogs (Schneider et al., 2018) to specifically evaluate the effect of FSS on Jag1-mediated Notch activation. In agreement with our previous data (Driessen et al., 2018), we found that FSS enhanced Jag1-mediated Notch activation. However, the levels of endothelial cell-induced activation of Notch in the reporter cells remained constant regardless of the magnitude of FSS stress tested (Fig. 1J). This contrasts with the FSS magnitude-dependent Jag1 expression levels (Fig. 1G-H) (Souilhol et al., 2022), indicating that FSS-regulated Jag1 levels may serve unique functions in endothelial cells unrelated to Notch activation. Even though endothelial-specific knock-out of Jag1 leads to embryonic lethality at E10.5 (High et al., 2008) Jag1 is considered a weak and redundant Notch activator in endothelial cells (Fernández-Chacón et al., 2021; Sprinzak and Blacklow, 2021; Sunshine et al., 2024). Our results further suggest an uncoupling of Jag1 and Notch activity, consistent with their distinct roles in various vascular processes.

### Jag1 inhibition impairs VEGFR2 and ERK activity

We next evaluated the involvement of Jag1 in endothelial mechanotransduction. FSS is known to be sensed by mechanoreceptors that activate downstream kinase signaling to regulate endothelial cell functions in the hemodynamic environment (Baeyens et al., 2016; Souilhol et al., 2020; Tamargo et al., 2023). We silenced Jag1 using siRNA and assessed the activity of FSS-responsive kinase activity in endothelial cells exposed to 0.8 Pa shear stress for 24 hrs in full media including growth factors (Fig. 2A). The phosphorylation levels of VEGFR2 were reduced in both static and flow conditions upon silencing Jag1 (Fig. 2B). Jag1 silencing also decreased the phosphorylation of ERK kinase induced by FSS (Fig. 2C) but caused no significant changes in the phosphorylation or total expression levels of AKT (fig. S3).

**Fig. 2.**
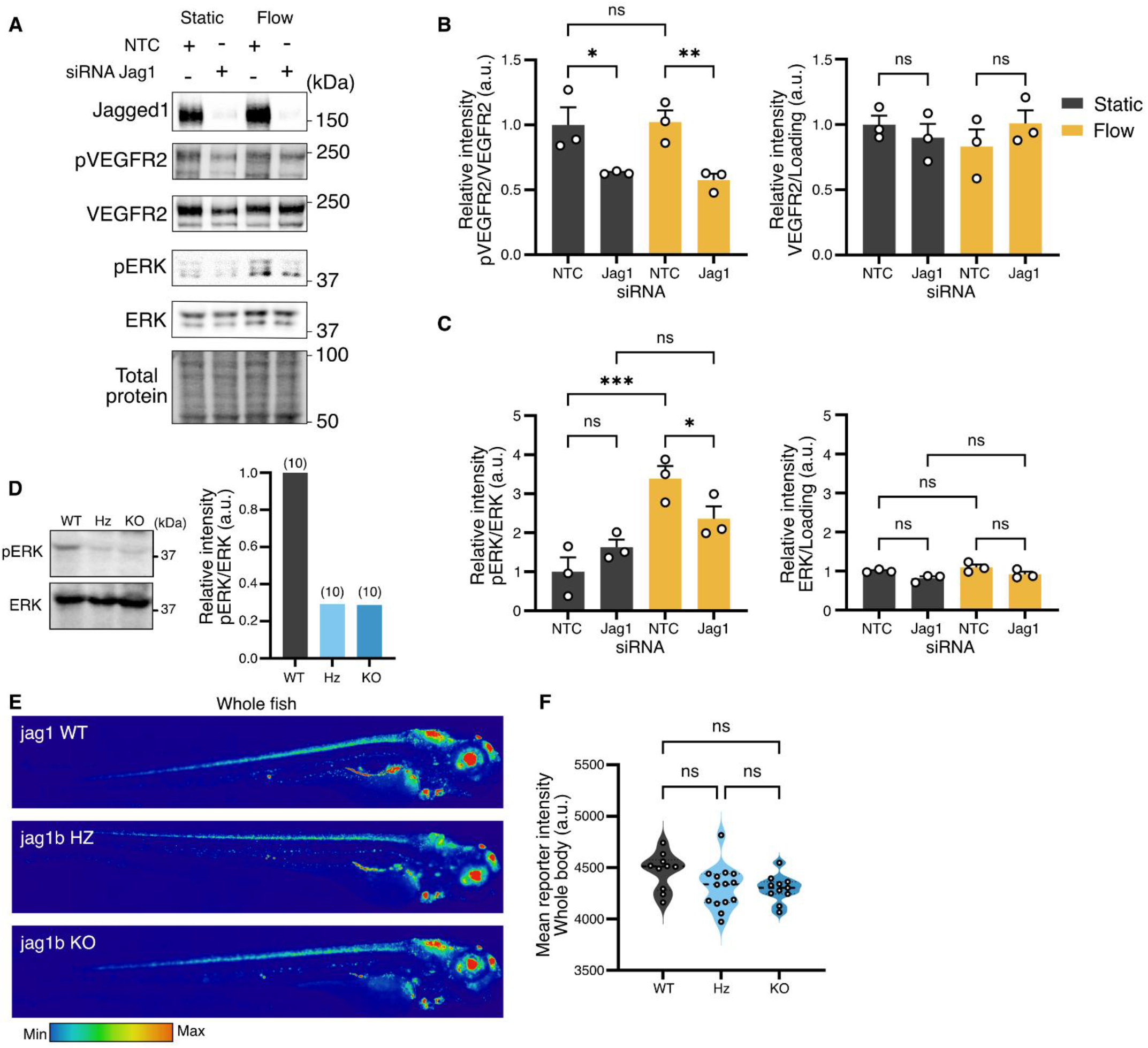
Jag1 is necessary for shear stress-linked kinase activity. (**A**) WB of the phosphorylation and protein expression levels of VEGFR2 and ERK in HUVECs exposed to 0.8 Pa of FSS in the orbital shaker. Cells were silenced for 48 hours with siRNA non-targeting control (NTC) or siRNA Jag1 before exposure to FSS for 24hours. Quantification of these experiments is found in (**B**) for VEGFR2 and (**C**) for ERK. (**D**) ERK activity was measured in protein lysates of WT (N=10), heterozygous (Hz) (N=10) and homozygous Jag1KO (KO) (N=10) embryos by WB. (**E**) Fluorescence images of Notch activity (TP1:H2B-mCherry) in TP1:H2B-mCherry (Jag1 (WT); TP1:H2B-mCherry/jag1bHz and TP1:H2B-mCherry/jag1bKO zebrafish at 4 dpf. Scale bar: 200 µm for the heart. Quantification of Notch activity in the the whole fish (**F**) was performed by measuring fluorescent intensity. P-values were obtained with GraphPad Prism as described in the methodology. Significance is indicated as: ns p > 0.05, * p < 0.05, ** p < 0.01, *** p < 0.001. Arbitrary units are indicated as (a.u.).

To test whether Jag1 also affected kinase signaling in vivo, we utilized stable Jag1b KO zebrafish (*Danio rerio*) embryos. The zebrafish cardiovascular system begings to develop during the first 24 hours post fertilization (hpf) and the onset of flow occurs 25-26 hpf (Isogai et al., 2001). Notch activity has been shown to be induced by flow in the vasculature at 2 days post fertilization (dpf) (Weijts et al., 2018). Zebrafish express two forms of Jag1: jag1a and jag1b. Whereas the extracellular and intracellular domains of jag1b show high similarity to human Jag1, the intracellular domain of jag1a only has 30% similarity with substantial variations in the PDZ-binding motifs (PDZBM) that mediate protein-protein interactions (fig. S4A). We analyzed kinase activity in lysates from WT, jag1b heterozygous, and jag1bKO zebrafish embryos at 4 dpf. The VEGFR2 antibody was incompatible with zebrafish, but ERK activity was significantly reduced in the heterozygote and jag1bKO compared to WT embryos (Fig. 2D). Since both canonical and non-canonical Jag1 signaling has been shown to affect ERK activity (Pelullo et al., 2019; Sanna et al., 2020), we crossed the Jag1bKO fish with the tg(*TP1:H2B-mCherry*) Notch reporter line (Ninov et al., 2012) to analyze Notch activity. We did not observe any significant changes in Notch signaling activity upon deletion of jag1b (Fig. 2E), measured by the fluorescence reporter intensity in the whole fish (Fig. 2F). The jag1bKO did show pericardial edema (fig. S4B-C) consistent with observations in zebrafish treated with kinase inhibitors (Anastasaki et al., 2012; Bolcome and Chan, 2010; Ma et al., 2022), although the exact underlining causes of this effect needs to be determined. Taken together, the data suggests that Jag1 may play a non-canonical role in regulating the activity of mechanosensitive kinases in endothelial cells.

### Physical stimulation of Jagged1 induces VEGFR2 activity

Since Jag1 depletion impaired kinase activity, we next examined if Jag1-mediated kinase activity could be induced by direct stimulation of Jag1. To this end, we first applied tensional force on Jag1 using 1 µm paramagnetic beads coated with an antibody that recognizes the extracellular domain of Jag1 (J1ECD ab) (Fig. 3A). A similar system has been used to evaluate the mechanosensitivity of proteins involved in FSS induced signaling (Mehta et al., 2020; Tzima et al., 2005). Antibody validation using Jag1 wild-type and knockout cells is presented in fig. S5. We exposed endothelial cells to the beads for 15 minutes, followed by an additional 15 minutes of exposure to a magnetic force field. VEGFR2 activation was increased in endothelial cells exposed to J1ECD ab-coated beads but not in those exposed to NECD ab-, rNECD-, or rIgG-coated beads (Fig. 3B). Exposure to J1ECD ab-coated beads also induced ERK activation (fig. S6A).

**Fig. 3.**
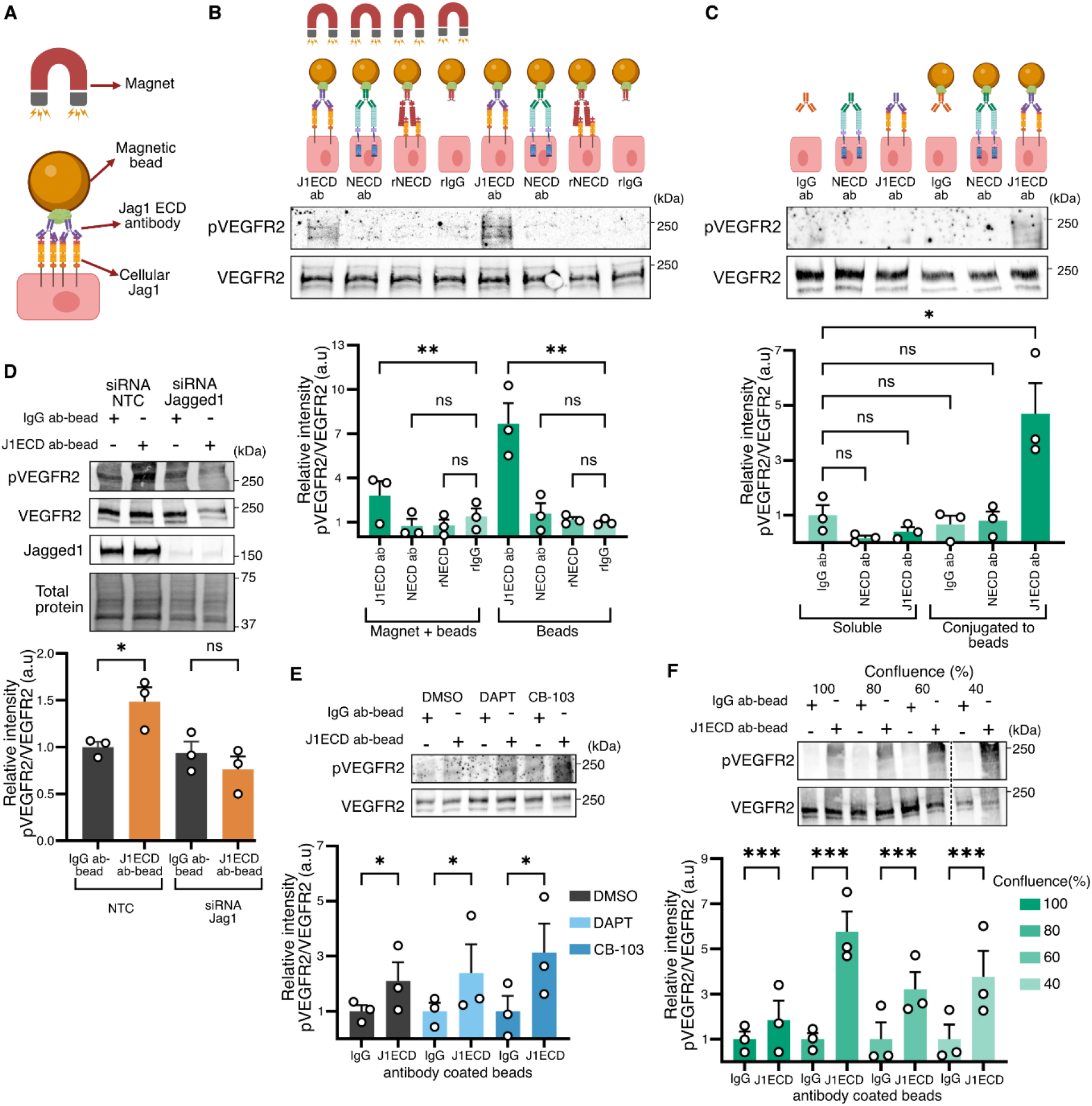
Mechanical stimulation of Jag1 activates shear stress-related kinases independently of Notch. (**A**) Schematic illustration (not to scale) of the magnetic bead experiment: HUVECs were incubated with protein A/G magnetic beads coated with Jag1 extracellular domain (ECD) antibodies, (J1ECD ab), Notch ECD antibodies (NECD ab), recombinant Notch ECD (rNECD), or recombinant IgG (rIgG) for 15 minutes and then exposed to a magnet or incubated with the cells for a total of 30 minutes. (**B**) WB and analysis (below) of VEGFR2 and phosphorylated VEGFR2 in HUVECs treated with magnetic beads coated with J1ECD ab, NECD ab, rNECD, or rIgG. VEGFR2 phosphorylation was induced only in cells incubated with magnetic beads coated with the Jag1ECD antibody. (**C**) WB analysis of VEGFR2 and phosphorylated VEGFR2 in HUVECs treated with soluble J1ECD ab, NECD ab, or IgG ab or conjugated to magnetic beads. J1ECD ab-coated beads but not soluble Jag1 antibodies induced VEGFR2 phosphorylation. (**D**) WB analysis of VEGFR2 and phosphorylated VEGFR2 in control and Jag1-silenced HUVECs stimulated by J1ECD ab- or IgG ab-coated beads. siRNA-mediated silencing of Jag1 inhibited phosphorylation of VEGFR2 induced by the J1ECD ab-coated beads. (**E**) WB analysis of phosphorylated and total VEGFR2 in HUVECs treated with J1ECD ab- or IgG ab-beads in the absence and presence of Notch pathway inhibitors. Pretreatment with Notch inhibitors DAPT and CB-103 did not reduce the Jag1-mediated VEGFR2 response. Values were normalized to the IgG ab-bead control. (**F**) WB analysis of phosphorylated and total VEGFR2 in HUVECs of different confluences treated with J1ECD ab- or IgG ab-coated beads. J1ECD ab-coated beads induced VEGFR2 activation did not decline with decreasing confluence in HUVECs. The values were normalized to the IgG ab-coated bead control for each confluence. The levels are presented as the mean of each replicate relative to their corresponding control+ SEM. HSC-70 and whole cell lysates were used as a loading control. P-values were obtained with GraphPad Prism as described in the methodology. Significance is indicated as: ns p > 0.05, * p < 0.05, ** p < 0.01, *** p < 0.001. Arbitrary units are indicated as (a.u.).

The presence of the magnetic field resulted in a minor increase in Jag1-induced VEGFR2 activation compared to those without it (Fig. 3B), indicating that mechanical force might not be necessary to induce VEGFR2 activity in response to Jag1 stimulation. However, exposing cells to soluble antibodies did not induce VEGFR2 activation, indicating that a physical component is necessary for Jag1-induced VEGFR2 activation (Fig. 3C). This finding is consistent with observations made for VE-Cadherin (Seo et al., 2016) and integrins (Arias-Salgado et al., 2003; Miyamoto et al., 1995; Plopper et al., 1995), where clustering induced by antibody-coupled microbeads, without further force, has been shown to induce signal activation. To further confirm the involvement of Jag1 in VEGFR2 activation, we silenced Jag1 in the endothelial cells. siRNA-mediated Jag1 knockdown prevented VEGFR2 phosphorylation by the J1ECD ab-coated beads (Fig. 3D).

While rNECD-coated beads were expected to interact and stimulate Jag1, exposing cells to these beads for 30 minutes was insufficient to produce a detectable increase in VEGFR2 phosphorylation (Fig. 3B-C). We also tested if cells plated on top of immobilized rNECD could activate phosphorylation of VEGFR2 and ERK (fig. S6B). Increase in phosphorylation was indeed detected but only after 6 hours of exposure (fig. S6C-6D). These findings align with previous reports, which show that beads coated with the recombinant Jag1 ligand induce Notch activation after days of incubation (De La Croix Ndong et al., 2018; Zohorsky et al., 2021). This longer time requirement may be due to the low affinity of the interaction between Jag1 ligands and Notch receptors compared to that of monoclonal antibodies.

To assess if Jag1-induced VEGFR2 activation required canonical Notch activity, we treated endothelial cells with Notch inhibitors. The γ-secretase inhibitor DAPT was used to prevent Notch cleavage and consequently inhibit canonical (Song et al., 2023) and non-canonical cortical Notch activation (Polacheck et al., 2017). The more recent inhibitor, CB-103, was used to inhibit Notch transcriptional activation (Lehal et al., 2020). We cultured the cells in the presence of the inhibitors for 24 hours before exposing them to beads, to ensure efficient Notch inhibition. Neither inhibition of Notch cleavage nor transcriptional activity prevented VEGFR2 activation induced by the J1ECD ab-coated beads (Fig. 3E). Since canonical Notch activity requires cell-cell contact, seeding the cells at lower confluences reduces Notch activity. We further demonstrated that lowering the confluence of the cultures did not decrease VEGFR2 activation (Fig. 3F). Taken together, the data indicates that Jag1-mediated VEGFR2 activation does not require canonical Notch activity.

### Jag1-mediated signaling requires Src activity

We next assessed if direct interaction between Jag1 and VEGFR2 was needed to induce VEGFR2 phosphorylation. Jag1 and VEGFR2 did not interact in co-immunoprecipitation assays (Co-IP) (Fig. 4A). It is known that VEGFR2 is activated by the VEGFR2/PECAM/VE-Cadherin (Tzima et al., 2005) and PlexinD1/NRP1 (Mehta et al., 2020) mechanosensitive complexes. To understand if proteins involved in these complexes were important for Jag1-mediated VEGFR2 activation, we used the COS-7 cell line (Fig. 4B), which lacks the proteins of these junctional complex sensors of FSS (Mehta et al., 2020; Tzima et al., 2005). To our surprise, VEGFR2 was activated by J1ECD ab-coated beads in the absence of any of the other proteins in the complexes (Fig. 4C), indicating an alternative mode of VEGFR2 activation. Jag1 signaling has been shown to intersect with Src kinase signaling (Miloudi et al., 2019; Small et al., 2001; Trifonova et al., 2004), and Src has been shown to operate both downstream and upstream of VEGFR2 (Aitken et al., 2023; Jin et al., 2003; Miller and Sewell-Loftin, 2022). We analyzed Src kinase activity in the COS-7 cells treated with the J1ECD ab-coated beads. Jag1 stimulation induced Src and VEGFR2 activity without any other protein of the mechanosensory complexes. Furthermore, Jag1 stimulation induced Src activation even in the absence of VEGFR2 (Fig. 4C), indicating that Src acts upstream of VEGFR2. We found that Src co-immunoprecipitated with Jag1, indicating a potential direct interaction (Fig. 4D), although the exact mechanisms of Jag1-mediated Src activation needs to be determined. To test if Src activity was necessary for VEGFR2 phosphorylation in endothelial cells, we stimulated HUVECs with J1ECD ab-coated beads in the presence and absence of the Src inhibitor Saracatinib. Inhibition of Src prevented VEGFR2 phosphorylation, demonstrating that Jag1-mediated VEGFR2 activation required Src activity (Fig. 4E).

**Fig. 4.**
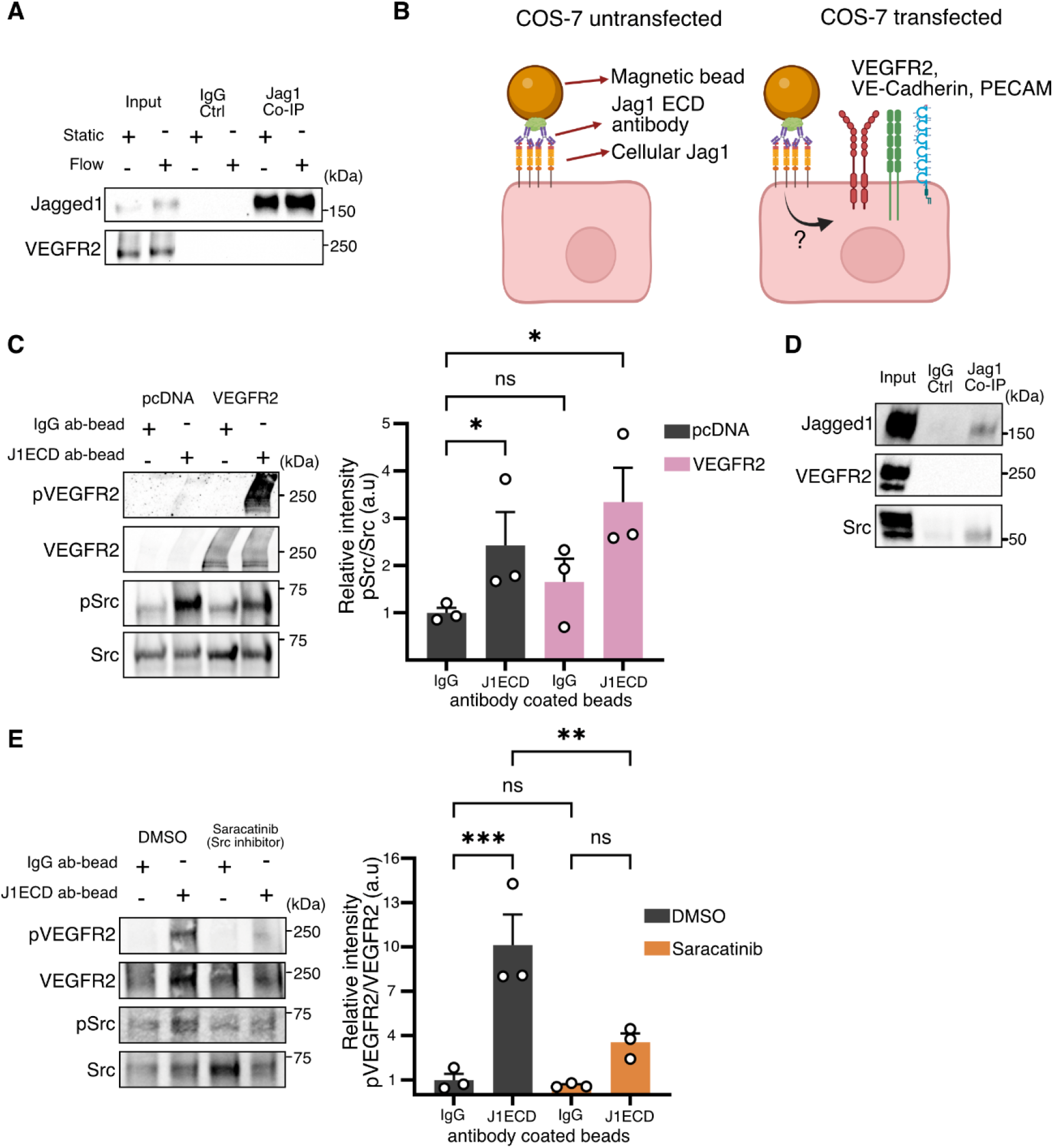
Jag1-mediated VEGFR2 activation requires Src. (**A**) Jag1 Co-Immunoprecipitation (Co-IP) analysis by WB, of VEGFR2 in HUVECs cultured in static conditions or exposed to flow (0.8 Pa FSS) using the orbital shaker system. No VEGFR2 was detected in Jag1 Co-IP samples. (**B**) Schematic of transfection experiment with COS-7 cells. (**C**) WB analysis of total and phosphorylated VEGFR2 and Src in Cos-7 stimulated with J1ECD ab- or IgG ab-coated beads after being transfected with plasmid control (pcDNA) or VEGFR2 expression plasmid. Stimulation of Cos-7 cells with J1ECD ab-coated beads induced Src and VEGFR2 activation. Src was activated in the absence of VEGFR2. The values were normalized to the IgG ab-bead control. (**D**) Jag1 Co-IP analysis by WB of VEGFR2 and Src in HUVECs cultured in static conditions. Src but not VEGFR2 are detected in Jag1 Co-IP samples. (**E**) WB analysis of phosphorylated and total VEGFR2 and Src in HUVECs stimulated with J1ECD ab- or IgG ab-coated beads in the presence and absence of the Src inhibitor Saracatinib. The values were normalized to the IgG bead control. The levels are presented as the mean of each replicate relative to their corresponding control + SEM. P-values were obtained with GraphPad Prism as described in the methodology. Significance is indicated as: ns p > 0.05, * p < 0.05, ** p < 0.01, *** p < 0.001. Arbitrary units are indicated as (a.u.).

## CONCLUSIONS

Taken together, our data describe a new non-canonical role of Jag1 in regulating the activity of kinases involved in endothelial mechanotransduction. Physical stimulation of Jag1 induces Src activity, which in turn is required for Jag1-mediated VEGFR2 activation. Since exposure to J1ECD-coated beads was sufficient to activate Src without requiring a magnetic force, but the beads were necessary, we hypothesize that the relocalization and clustering of Jag1 and Src interaction may trigger Src activation, in a similar manner as what has been observed with integrins (Arias-Salgado et al., 2003; Miyamoto et al., 1995; Plopper et al., 1995). The clustering induced by the antibody-coated beads might be analogous to the relocalization and clustering of Jag1 induced by FSS we observe in endothelial cells exposed to shear stress.

Although Jag1-mediated VEGFR2 activation appears to be mediated through Src, the underlying mechanism remains unclear. Other groups have previously shown that the cleaved extracellular domain of Jag1 is capable of activating Src in the vasculature by modulating both canonical and non-canonical Notch signaling pathways (Miloudi et al., 2019). However, we demonstrated that Jag1-mediated VEGFR2 activation does not require Notch activity as VEGFR2 can be activated in the presence of the γ-secretase inhibitor DAPT that inhibits both canonical and non-canonical Notch activity (Polacheck et al., 2017; Song et al., 2023).

Our data could help explain some of the discrepancies between the effects of deleting Jag1 and Notch1 in the endothelium. We observed that Jag1 re-localizes in response to FSS polarizing downstream of the flow direction. Notch has previously been shown to also polarize in the direction of flow. Since cis-interactions with Jag1 inhibit Notch activation (Giese et al., 2025; Mack et al., 2017), our data could help explain the contrasting effects of Notch and Jag1 in atherosclerosis (Mack et al., 2017; Souilhol et al., 2022). The pro-atherogenic role of Jag1 may be related in part to cis-inhibition of Notch by Jag1 in regions of low oscillatory flow or to Notch-independent functions in kinase signaling. Additionally, while we found Jag1 to be a positive regulator of ERK activity, other groups have shown that Notch is a negative regulator of endothelial ERK activation in mouse and zebrafish (Pontes-Quero et al., 2019; Shin et al., 2016). On the same line, unlike Notch1 (Mack et al., 2017), Jag1 and ERK are pro-atherogenic (Mei et al., 2012; Souilhol et al., 2022; Wang et al., 2025). More research will be needed to fully clarify the interplay of Jag1 and Notch in vascular mechanotransduction.

## MATERIALS AND METHODS

### Cell culture

Pooled human umbilical vein endothelial cells (HUVEC) (PromoCell) were cultured in Endothelial Cell Growth Medium 2 (PromoCell) completed with Endothelial Cell Growth Medium 2 SupplementMix (PromoCell). Human Aortic Endothelial Cells (HAoEC) (PromoCell) were cultured in Endothelial Cell Growth Medium MV (PromoCell) completed with Growth Medium MV SupplementMix (PromoCell). Thawing and expansion of primary endothelial cells were executed according to the PromoCell Instruction Manual, and all experiments, unless stated otherwise, were done using fully confluent cells at passages 5-6. COS-7 cells were cultured in DMEM (Sigma) supplemented with 10% FBS, 100 U/ml penicillin, and 100 µg/ml streptomycin. All cells were seeded and expanded on tissue culture polystyrene (TCPS) plates and maintained at 37 °C, 5% CO2.

### Experimental animals

Jagged1b (b1105) (Zuniga et al., 2010) was generously provided by the Crump Lab at the University of Southern California, Keck School of Medicine. All the experiments were carried out at Turku Bioscience Center Zebrafish Core under license ESAVI/31414/2020 and ESAVI/44584/2023 granted by The Regional Administration Office of Southern Finland. jag1b +/-strain and TP1:H2B-mCherry; jag1b+/-strain were created in Turku by crossing jag1b +/-together with tg(*TP1:H2B-mCherry*) strain (Ninov et al., 2012) generously provided by the Ninov Lab at the Center for Regenerative Therapies Dresden (CRTD) at TUD Dresden University of Technology. Embryos used for imaging were kept in E3 media (5 mM NaCl, 0.17 mM KCl, 0.33 mM CaCl2, 0.33 mM MgSO4) supplemented with 30 mg/ml of 1-phenyl 2-thiourea (PTU) in 28.5°C. Imaging was carried out using Zeiss AxioZoom V.16 stereomicroscope with 1.0x Plan ApoZ (NA 0.125) objective and Hamamatsu sCMOS ORCA-Flash4.0 LT. During the imaging, embryos were anesthetized with Tricaine 160 mg/ml. Notch activity was measured using ImageJ FIJI image analysis software. For western blot (WB) analysis, embryos were genotyped and then pooled accordingly. Embryos were lysed with 3x Laemmli buffer (30 % Glycerol, 3 % SDS, 0.1875 M Tris-HCl pH 6.8, 0.015 % bromophenol blue, 3% β-mercaptoethanol) and boiled 15 minutes at 95°C.

For genotyping, the genomic DNA was extracted from each embryo via alkaline lysis using 50 mM NaOH and boiling the samples for 15 minutes at 95°C. Samples were vortexed every 5 minutes and at the end centrifuged 5 minutes, 14 000 relative centrifugal force (rcf) at 4°C. DreamTaq DNA polymerase (Thermo Scientific) was used in gene amplification according to the manufacturer’s instructions followed by BseGI (BtsCI) (Thermo Fisher Scientific)restriction enzyme treatment. Following primers were used in gene amplification: Forward primer: GTACCAAATCCGGGTGACCT, Reverse primer: GTGGCTTTTTGGGTCATTATCA. Samples were analyzed from 2.5% Agarose gel. BseGI treatment resulted fragments listed below.

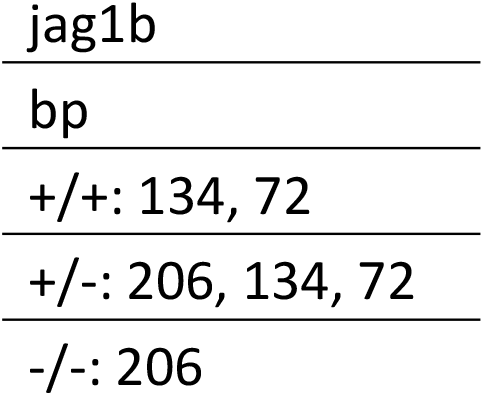

### Shear stress experiments

For imaging experiments, HUVECs and HAoECs were seeded into 6-channel slides (Ibidi®, 80606 coated with 100 µg / ml Bovine Type I Collagen solution (Advanced BioMatrix Inc., 5010) with 1 × 10^5^ cells added per channel. Media was changed twice a day. Cells were allowed to attach and form a monolayer overnight, after which they were subjected to shear stress or maintained as static control. Flow over cells was achieved by assembling a perfusion set consisting of the 6-channel slide connected to a glass bottle with 25 ml of respective culture media via 1.6 mm inner diameter silicone tubing (Ibidi®, 10842) and attached to a REGLO Analog peristaltic pump (Ismatec) via three-stop-tubing, creating a loop for media recirculation during the experiment. Utilizing a three-port screw cap on the glass bottle, a 22 µm pore diameter syringe filter was attached to the bottle to allow air pressure equilibration in the system. At the start of a shear stress experiment, the flow rate of the system was increased in a step-wise manner over one hour by changing the pump rpm, starting from 0 rpm, followed by 22, 66, and 99 rpm steps, 99 rpm equaling ∼1 Pa of laminar and continuous FSS. Cells were exposed to flow for 24 to 25 hours at ∼1 Pa. The media was preconditioned to the temperature, humidity and CO2 conditions of the experiment by putting it in a glass bottle inside the incubator overnight before the assembly of the perfusion set.

For gene expression experiments, HUVECs were seeded at 10^6^ cells/ml into collagen IV-coated 1-channel slides for gene expression analysis and 6-channel slides (Ibidi®, 80172) for immunocytochemistry or the reporter cell assay. After one day of culture, the endothelial cells were subjected to FSS using the Ibidi® pump system (Ibidi®) or kept in culture as a static control. The standard perfusion sets were adjusted to gain a stable flow speed and pressure. Resistance tubing (0.5 mm inner diameter) was added after the channel slides to stabilize the fluctuations in flow speed. The cells were exposed to 0.5, 1, or 2 Pa FSS for 24 hours. Flow speeds were constantly measured to ensure correct flow speed (ME2PXL flow sensor, Transonic Systems Inc.).

For protein and phosphorylation analysis, cells were seeded in 6-well plates and grown to full confluency. Before flowing, in the beginning of the experiment, the media was changed to ensure an initial volume of 3mL per well and the plate was placed on a CO2-resistant orbital shaker. A computational fluid dynamics simulation was performed as previously described (Driessen et al., 2020) for a 19mm shaking orbital diameter. The simulation values were used to select the speed of the orbital shaker, and in all cases, a non-flowed plate (Static) was used as a control. Unless stated otherwise, plates were collected after 24 hours and placed on ice, and approximately 64% of the outermost area of the well was collected.

### Immunocytochemistry

After the shear stress experiment, cells in the 6-channel Ibidi® slides were washed two to three times with 37°C modified (without CaCl_2_, MgCl_2_) Dulbecco’s Phosphate-Buffered Saline (DPBS) and fixed in 37°C 4% paraformaldehyde in DPBS for 10 minutes, followed by another three washes with modified DPBS. Slides with fixed cells were stored for up to 16 days at a temperature of 4°C in the dark. Cells were permeabilized in 0.2% Triton X-100 in DPBS for 5 and 10 minutes at room temperature (RT), after which they were washed once with modified DPBS. Blocking of the cells was done by incubation in 1% BSA in modified DPBS at RT for 30 minutes, followed by three washes with modified DPBS. Cells were incubated with Jag1 (Cell Signaling Technology; 2620; 1:100) and CD31 (Invitrogen; 37-0700; 1:100) monoclonal primary antibodies in the solution of 3% BSA and 0.05% Triton X-100 in DPBS overnight at 4°C. Cell was washed three times with modified DPBS and incubated with secondary antibodies (Donkey anti-Mouse IgG Alexa Fluor 488; A-21202; 1:500, Goat anti-Rabbit IgG Alexa Fluor 555; A-21428, 1:500) and DAPI (Sigma-Aldrich, D9542) in 3% BSA and 0.05% Triton X-100 in PBS for 1 hour at RT in the dark. The staining solution was removed, and cells were washed three times in RT-modified DPBS and imaged with the channels filled with PBS.

Imaging was conducted with a Zeiss LSM 880 Airyscan confocal with an Axio Observer.Z1 microscope using ZEN 2.3 SP1 black edition acquisition software. The objective used was 63x Zeiss C Plan-Apochromat Oil DIC M27 with 1.4 aperture. The pinhole was set to give the same optical section thickness for all channels. DAPI was excited with a diode at a wavelength of 405 nm and acquired at an emission window of 410-514 nm with a PMT with the pinhole set to 3.85 airy units (AU). Alexa Fluor 488 was excited with Argon laser at a wavelength of 488 nm and acquired at an emission window of 490 – 579 nm with GaAsP detector with pinhole set to 3.10 AU. Alexa Fluor 555 was excited with HeNe laser at a wavelength of 543 and acquired at an emission window of 556 – 648 with a cooled PMT with the pinhole set to 2.69 AU. Channels were acquired sequentially and with unidirectional scanning, 4 times line averaging, and pixel dwell time of 1.02 µs at a bit depth of 8 bits. All images were acquired with 1×1 binning, and the XYZ pixel dimensions varied across experiments within 85 – 132 nm in X and Y and 632-1090 nm in Z.

PolarityJaM (Giese et al., 2025) was used to quantitatively measure the polarization of Jag1 from confocal images. To pre-process the confocal 3D images for PolarityJaM, two channel 2D images were generated from the confocal 3D image stacks by maximum intensity projection of the Jag1 channel and by picking optical section corresponding to the base of the cell from PECAM-1 channel. In PolarityJaM, PECAM-1 channel was used as the junction channel to generate cell masks and signal polarity was measured from Jag1 channel. PolarityJaM parameters used in the analysis will be available in the repository upon acceptance for publication.

### Co-immunoprecipitations and Western blots (WB)

Cells were collected in lysis buffer (50 mM Tris, pH 7.5, 150 mM NaCl, 1% Triton X-100, 0.1% SDS) with protease (Complete™ Protease Inhibitor Cocktail, Merck) and phosphatase inhibitor cocktail (Pierce™ Phosphatase Inhibitor Mini Tablets, ThermoFisher). Lysates were collected and kept on ice during sonication before being centrifuged for 10 minutes at 14 000 rcf at 4°C. The supernatant was used for the input control and the subsequent processing steps. Beads were washed five times with washing buffer (50 mM Tris-HCl (pH 7.5); 250 mM NaCl; 0,1% NP-40) before use. Lysates were pre-cleared using 5 µL of washed magnetic beads per sample. After pre-clearing, samples were incubated overnight at 4 °C with antibody (Jag1 28H8 1:50, VEGFR2 55B11 1:100 or IgG (DA1E); all from Cell Signaling Technologies (CST)). Samples were then incubated at RT for 1.5 hours with 16.5 µl of washed beads per sample. Finally, beads were washed three times with a washing buffer and, one final time, with MilliQH_2_O before being diluted in Laemmli sample buffer for WB.

Proteins were separated by SDS-PAGE and transferred to a Protran nitrocellulose membrane (GE Healthcare Life Sciences) using a wet transfer apparatus (Amersham Bioscience). The membranes were blocked with 5% nonfat dry milk in TBS at RT for 0.5-1 hour. Primary antibody incubation was done overnight at 4 °C in constant agitation. Membranes were then incubated in secondary antibody for 1 hour at RT (1:10000). Proteins were acquired using an iBright Imaging System (ThermoFisher) after incubation with SuperSignal West Pico PLUS Enhanced chemiluminescence substrate (ThermoFisher). The following antibodies were used: Jag1 28H8, VEGFR2 55B11, phospho-Tyr1175 VEGFR2 2478, p44/42 MAPK (Erk1/2) 9102S, phospho-Y204-Erk1 / phospho-Y187-Erk2 5726S, Akt 9272, phospho-Ser473 Akt 4060S, CD31 (PECAM-1) D8V9E, VE-Cadherin D87F2, Src 2108S and Phospho-Src Family (Tyr416) D49G4 all purchased from CST. HSC70 (ADI-SPA-815-D, Enzo) or Revert 700 Total Protein Stain (LI-COR) was used for loading control. The densitometry level were obtained using the image analysis software ImageJ FIJI.

### Magnetic bead experiments

Experiments were performed using 24-well plates. 1.0 µm Protein A/G Magnetic Beads (ThermoFisher) were washed with TBS and coated for 1.5 hour at RT with antibodies (Jag1 1C4, IgG DA1E from CST; Notch 2 ECD MA5-24274 from ThermoFisher) or recombinant peptides (Recombinant Human Notch-1 Fc chimera protein and IgG-Fc chimera protein from R&D systems). Cells were incubated with beads at 37°C, 5% CO_2_ for 15 minutes before exposure to a magnetic field using permanent magnets for another 15 minutes or were incubated for 30 minutes before being lysed and collected for WB analysis.

### Pharmacological inhibitions

HUVECs were treated with the γ-secretase inhibitor 1N-[N-(3,5-difluorophenacetyl)-l-alanyl]-S-phenylglycine t-butyl ester (DAPT) to inhibit Notch cleavage or CB-103 to inhibit Notch-induced transcriptional activation, both at 25 µM. To inhibit Src, HUVECs were incubated with the inhibitor Saracatinib at 10 µM. All inhibitors were added in fresh media, 24 hours before experiments were performed. Inhibitors were dissolved in DMSO, and control experiments were performed in the presence of the same amount of DMSO as vehicle control.

### Gene expression

RNA was isolated with the Qiagen RNeasy kit. The β-mercaptoethanol – RLT buffer mixture was directly added to the cells in the 1-channel slides. The synthesis of cDNA was performed with M-MLV reverse transcriptase (Invitrogen). For six reference genes tested GAPDH was the most stably expressed, as analyzed with GeNorm130. The PCR protocol consisted of 3 minutes at 95°C, followed by 40 cycles of 20 s at 95°C, 20 s at 60°C and 30 s at 72°C. Data were analyzed using the ΔΔCt method. The primers used can be found below:

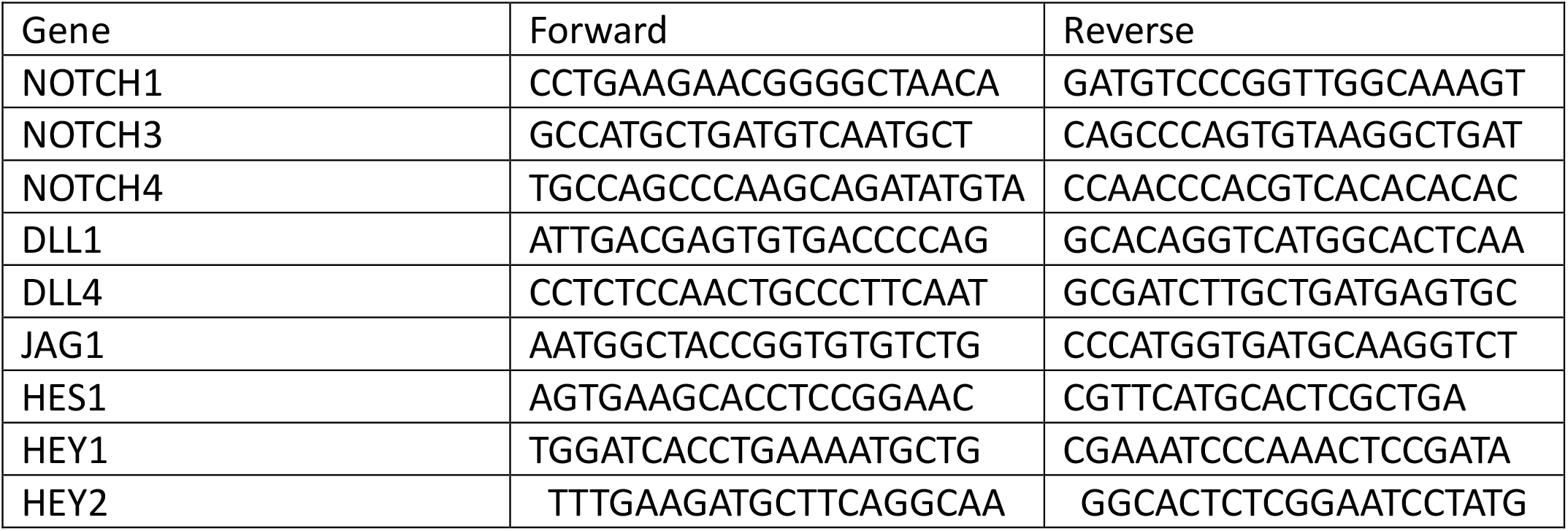

### Notch reporter assay

Notch1 overexpressing HEK293T cells were transfected with 12xCSL-luciferase223 or GFP as a transfection control using polyethyleneimine (PEI), 1 mg/ml. For transfection, DNA and PEI were mixed at 1:2 weight ratio in plain medium. After 5-minute incubation, the DNA-PEI mixture was added to the HEK293T cell culture. Next day, the cells were washed with PBS and used for the reporter assay. After flow experiments, endothelial cells were washed twice with PBS. Transfected HEK293T cells were seeded directly on top of endothelial cells (10^6^ cells/ml, 50 μl/channel) and cultured in EGM2 medium for 24 hours. In half of the channels (3), the cells were cocultured in the presence of 50 μM 6-alkynyl-fucose (Peptides International). In the other channels, the cells were cultured in an equal amount of DMSO as a control. Notch activity was assessed by measuring luciferase activity from lysed cells using Luciferase Assay from Promega. Biotek Synergy plate reader was used in signal detection.

### Statistical analysis

P-values were obtained after parametric tests were conducted. For two group comparisons a two-tailed Student’s T-test was performed. When three or more comparisons or more than one independent variable were present, two-way ANOVA were performed to assess main effects, with post-hoc testing. Dunnet’s post-hoc test was used when groups were compared with normalizing control. Tukey’s post-hoc test was used when all groups where compare with each other. Bonferroni correction was performed when four or more preselected comparisons against controls were made. For four or less comparisons Fishers LSD test was used as the post-hoc test. All error bars represent the SEM. The graphs were made using an established color-blind friendly palette (Wong, 2011). Graph and statistical analysis were performed using the statistical software GraphPad Prism 10.

## Supplementary Materials

fig. S1. Jag1 polarization in the direction of Flow in HUVECs.

fig. S2. Notch signaling gene expression at different magnitudes of FSS. fig. S3. Effect of Jag1 silencing on AKT activation.

fig. S4. Sequence comparison of Jag1 orthologs and pericardial edema in jag1b KO zebrafish.

fig. S5. Validation of Jag1 ICD and ECD antibodies.

fig. S6. Jag1-ab beads and immobilized Notch ECD induces VEGFR2 and ERK activation.

## Acknowledgments

The authors thank the ERC, the Academy of Finland, InFLAMES Research Flagship Center and Åbo Akademi University for their financial support; The Swedish Cultural Foundation in Finland and Instrumentarium Science Foundation for financially supporting the work of FSR; J. G. Crump (University of Southern California, Keck School of Medicine) for generously providing us with Jagged1b (b1105) zebrafish embryos; N. Ninov (Center for Regenerative Therapies Dresden (CRTD) at TUD Dresden University of Technology) for kindly providing us with TP1:H2B-mCherry zebrafish embryos; Cell Imaging and Cytometry Core (Turku Bioscience Centre and Biocenter Finland) for providing training and imaging facilities. I. Patero and the Zebrafish Core (Turku Bioscience Centre) for providing training and the facilities for the in vivo experiments on zebrafish; E. Långbacka, J. Chenglim Liu and A. Viitala (Åbo Akademi University) for their technical support. Figures were created with BioRender.com.

## Funding

This project has received funding from the following sources:

European Research Council (ERC) and the European Union’s Horizon 2020 research and innovation program grant agreement No 771168 (ForceMorph)

Research Council of Finland, decision number #316882 (SPACE)

Research Council of Finland, decision number #330411 (SignalSheets)

Research Council of Finland, decision number #336355 (Solutions for Health at Åbo Akademi University)

Research Council of Finland, decision number #337531 and #357911 (InFLAMES Flagship Program)

Åbo Akademi University Foundation’s Centers of Excellence in Cellular Mechanostasis (CellMech) and Bioelectronic Activation of Cell Functions (BACE)

The work performed by FSR has been partially funded by personal grants from The Swedish Cultural Foundation in Finland, Instrumentarium Science Foundation and Magnus Ehrnrooth Foundation.

## Author contributions

Conceptualization: FSR and CMS

Methodology: RCHD, FZ and OMJAS

Investigation: FSR, NV, EK, RCHD, FZ and OMJAS

Formal analysis: FSR, NV, EK, RCHD, FZ, OMJAS and CMS

Resources: CVCB and CMS

Data curation: OMJAS and FSR

Writing – original draft: FSR and CMS

Writing – review & editing: FSR, NV, EK, RCHD, FZ, CVCB, OMJAS and CMS

Visualization: FSR

Supervision: CVCB, OMJAS and CMS

Project administration: FSR and CMS

Funding acquisition: CVCB and CMS

## Data and materials availability

All data, code, and materials used in the analysis will be available in an open repository once the final form of the manuscript is approved for publication. All other data are available in the main text or the supplementary materials.

