## Supplementary figures for "A non-canonical role for Jagged1 in endothelial mechanotransduction"

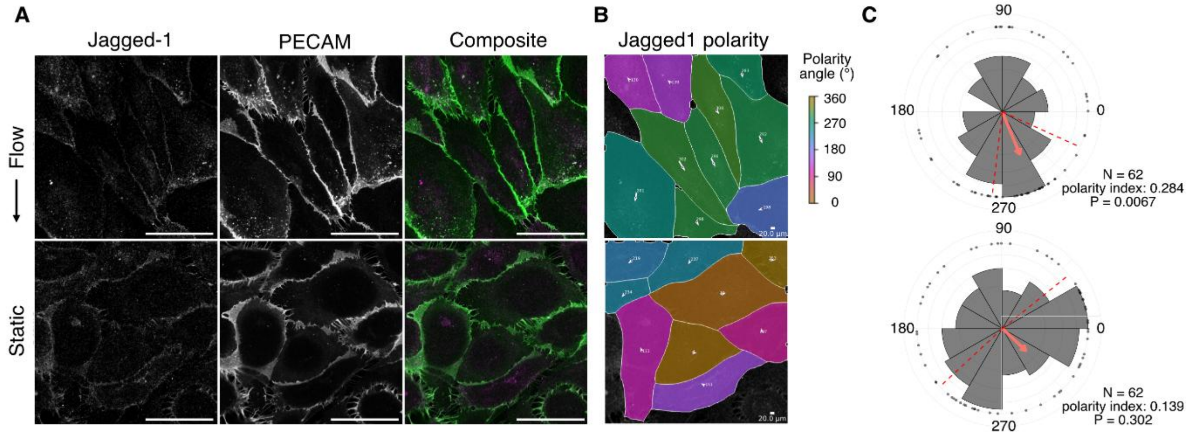


**fig. S1. Jag1 polarization in the direction of Flow in HUVECs.** (**A**) Confocal microscopy images of HUVECs exposed to ~1 Pa laminar and continuous FSS for 24 hours in Ibidi® microfluidic chips. Jag1 (magenta) demonstrated polarized localization in the direction of flow. PECAM/CD31 (green) was used to denote cell junctions. (**B**) Analysis of Jag1 polarization using PolarityJaM Python API (Giese et al., 2025). Mask color represents polarity angle. Vector size and orientation represent magnitude and angle of the polarization. (**C**) Merged graph and analyzes of three independent experiments in PolarityJaM web app. Vector represents the circular mean direction, dotted lines represent the 95% confidence interval. Rayleigh test of uniformity was used to analyze statistical significance.

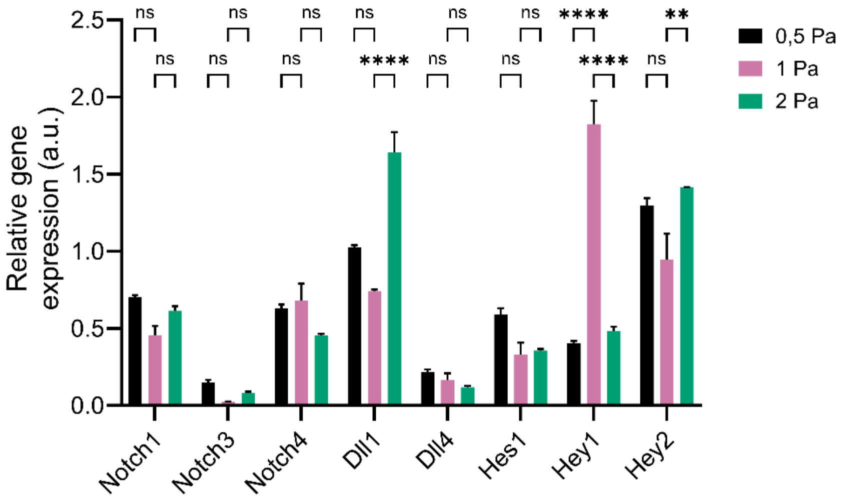


**fig. S2. Notch signaling gene expression at different magnitudes of FSS.** Q-PCR analyses of expression of Notch ligands and receptors in HUVECs exposed to 0.5, 1.0, and 2.0 Pa of laminar and continuous FSS for 24 hours in Ibidi® Chips. The experiment was performed three times with three technical replicates within each experiment. P-values were obtained with GraphPad Prism as described in the methodology. Significance is indicated as: ns p > 0.05 * p < 0.05, ** p < 0.01, *** p < 0.001. Arbitrary units are indicated as (a.u.)

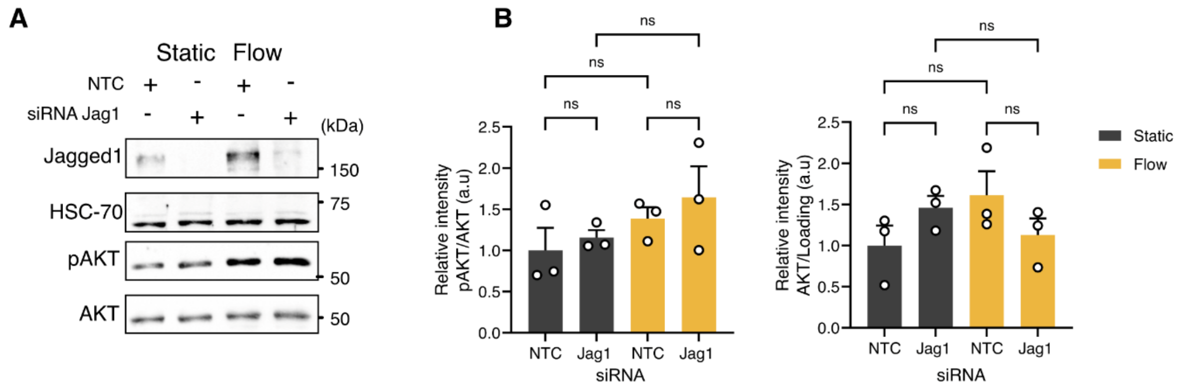


**fig. S3. Effect of Jag1 silencing on AKT activation** (**A**) AKT expression and activation in HUVECs exposed to 0.8 Pa of FSS in the orbital shaker. Jag1 was silenced by siRNA. The cells were incubated for 48 hours with siRNA non-targeting control (NTC) or siRNA Jag1 (Jag1) before exposure to FSS for 24 hours. (**B**) WB quantifications. The levels are presented as the mean of each replicate relative to their corresponding control + standard error of the mean (SEM). HSC-70 was used as a loading control. P-values were obtained with GraphPad Prism as described in the methodology. Significance is indicated as: ns p > 0.05 * p < 0.05, ** p < 0.01, *** p < 0.001. Arbitrary units are indicated as (a.u.)

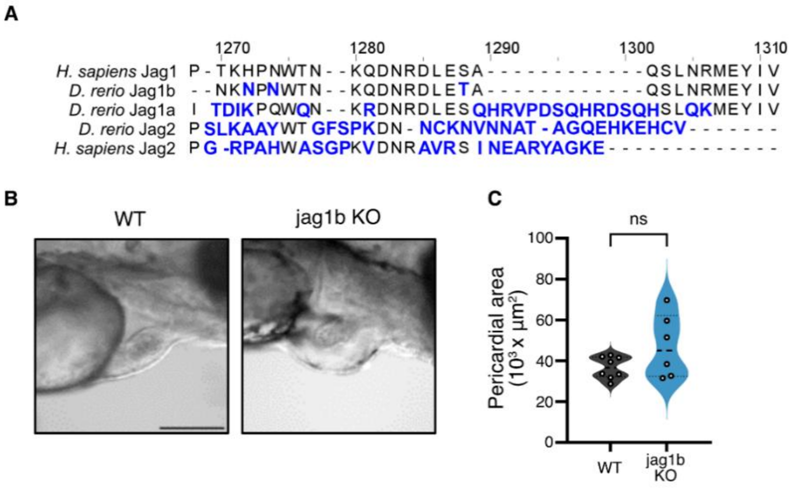


**fig. S4. Sequence comparison of Jag1 orthologs and pericardial edema in jag1b KO zebrafish.** (**A**) Zebrafish (D. rerio) express two orthologs of Jag1, Jag1a and Jag1b. Jag1b shares the human sequence with the last ten residues at the last N-terminal, including the PDZ-binding motif. Sequences were obtained from UniProt, and the multiple alignment analysis was performed in Clustal Omega. (**B**) Representative microscopy images of the developing heart in WT and Jag1b KO zebrafish at 4 dpf. jag1bKO zebrafish show signs of Pericardial edema. Scale bar: 200 µm. (**C**). Quantification of the pericardial area in Jag1WT and jag1bKO zebrafish at 4dpf. P-values were obtained with GraphPad Prism as described in the methodology. Significance is indicated as: ns p > 0.05, * p < 0.05, ** p < 0.01, *** p < 0.001.

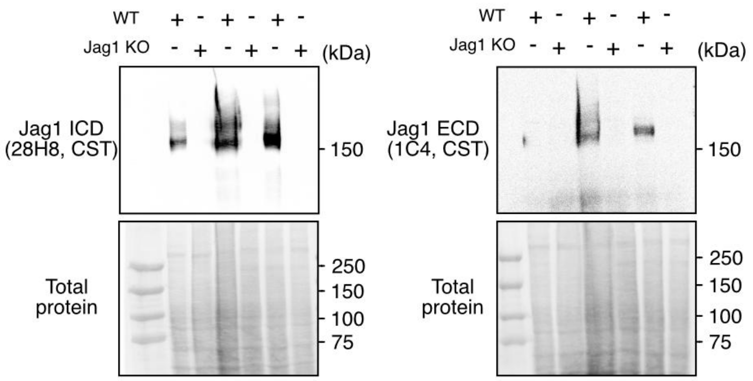


**fig. S5. Validation of Jag1 ICD and ECD antibodies.** Western blot of Jag1 knockout (KO) or WT BON1 cells with the following antibodies: Jag1 28H8 antibody from CST and Jag1 antibody 1C4 from CST, which recognizes Jag's extracellular domain (ECD). Representative blots from three independent experiments are shown. Both antibodies recognize Jag1 full length (180 kDa) specifically. We observe an unexpected unspecific signal around 50 kDa with the Jag1 1C4 antibody in both WT and Jag1KO. A similar pattern is observed in the total protein stain, indicating a potential bleed-through due to the weaker signal from the Jag1 1C4 antibody compared to that of the Jag1 28H8 antibody.

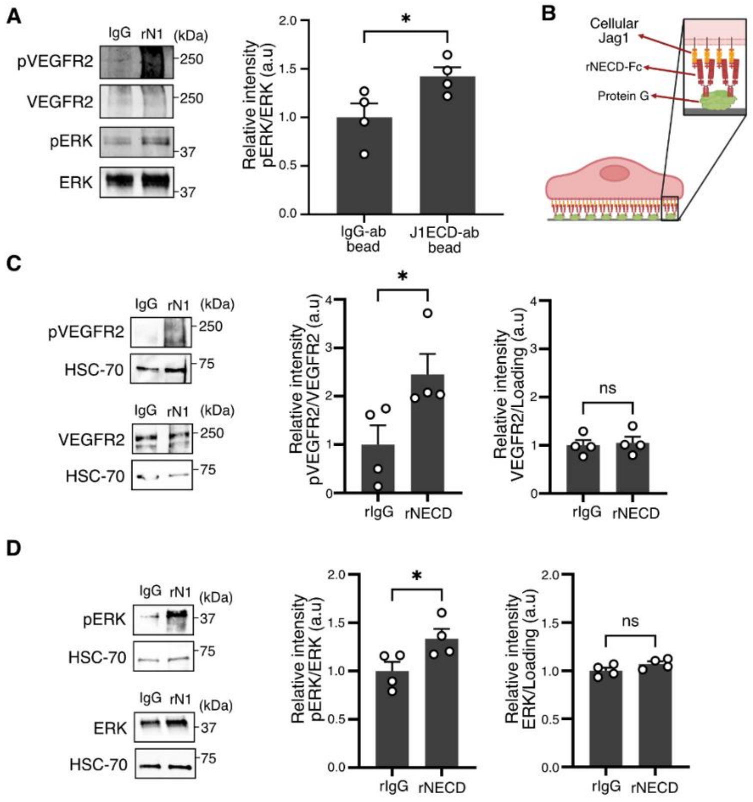


**fig. S6. J1ECD ab-beads and immobilized Notch ECD induces VEGFR2 and ERK activation.** (**A**) WB analysis of phosphorylated and total ERK levels in HUVECs incubated with protein A/G magnetic beads coated with Jag1 extracellular domain antibodies, (J1ECD-ab) or IgG antibody control (IgG-ab bead). (**B**) Schematic illustration of experimental setup with immobilized recombinant Notch extracellular peptides (rNECD): HUVECs were cultured for six hours on top of rIgG or rNECD immobilized to protein G-coated culture dishes. (**C**) WB analysis of phosphorylated and total VEGFR2 and (**D**) ERK levels in HUVECs cultured on rIgG or rNECD. The levels are presented as the mean of each replicate relative to their corresponding control+ SEM. HSC-70 and whole cell lysates were used as a loading control. P-values were obtained with GraphPad Prism as described in the methodology. Significance is indicated as: ns $p > 0.05$, * $p < 0.05$, ** $p < 0.01$, *** $p < 0.001$. Arbitrary units are indicated as (a.u.).
